# Optimal Cannabinoid-Terpene Combination Ratios Suppress Mutagenicity of Gastric Reflux in Normal and Metaplastic Esophageal Cells

**DOI:** 10.1101/2025.09.23.678062

**Authors:** Aaron Goldman, Gabriel Gonzalez, Svetlana A. Karpova, Leutz Buon, Masood A. Shammas, Hiroshi Mashimo, Markus H. Frank, Natasha Y. Frank

## Abstract

**Background:** Esophageal adenocarcinoma (EAC) frequently arises from chronic exposure to acid and bile reflux, with secondary bile acids, such as deoxycholic acid (DCA), contributing to its pathogenesis through mechanisms involving reactive oxygen species (ROS), oxidative DNA damage, and resistance to apoptosis. The human endocannabinoid system (ECS) regulates diverse anti-inflammatory, antioxidant, and analgesic pathways implicated in disease modulation. Despite its therapeutic promise, effective pharmacological activation of the ECS remains challenging.

**Objectives:** This study aimed to evaluate whether specific cannabinoid-terpene combinations targeting the ECS could attenuate the mutagenic and cytotoxic effects of bile acid–induced stress in esophageal cell models. Additionally, we assessed the clinical significance of ECS-related protein receptors in the progression of EAC.

**Design:** *In vitro* experimental models combined with clinical samples analyses.

**Methods:** We utilized *in vitro* models, including human esophageal epithelial cell lines exposed to DCA and a Barrett’s esophagus gastroesophageal reflux (GER) model subjected to low pH and a bile acid cocktail. Patient-derived samples were analyzed to investigate the clinical association of ECS pathway markers with EAC progression. Experimental models were treated with varying ratios of phyto-cannabinoids and terpenes. Endpoints included assessment of DNA damage, mitochondrial membrane potential, and ROS production to identify optimal compound combinations. Expression of ECS-related protein receptors was evaluated in clinical samples to elucidate their role in EAC development.

**Results:** A 1:5 ratio of cannabigerol (CBG) to Phytol (Phy) was found to significantly reduce DCA-induced DNA damage, preserve mitochondrial membrane potential, and decrease ROS levels. This combination also enhanced apoptosis in damaged cells and diminished mutagenicity. Analysis of patient samples revealed that the expression of the ECS-associated receptor protein CB1 correlated with EAC progression, suggesting a broader clinical role for ECS modulation in cancer prevention.

**Conclusion:** Modulation of the ECS through carefully selected cannabinoid-terpene ratios can mitigate bile acid–induced esophageal damage and may reduce carcinogenic progression. These findings support further *in vivo* investigations and raise the possibility of expanding cannabinoid-terpene therapeutics to other conditions involving similar pathogenic processes.

## Introduction

Esophageal adenocarcinoma (EAC) is a highly aggressive malignancy associated with Barrett’s Esophagus (BE) dysplasia and metaplasia ^1-3^, conditions affiliated with chronic exposure to acid-biliary reflux and gastroesophageal reflux disorder (GERD) ^4-6^. Bile acids, such as deoxycholic acid (DCA), enter the esophagus during episodes of reflux and are thought to promote cancer development ^7^. Patients with GERD and BE show high concentrations of DCA in their refluxate. DCA has cytotoxic effects and can induce DNA damage through a process that involves the induction of reactive oxygen species (ROS)^8-10^ and disruption of lysosomal integrity, which can drive ionic perturbations ^11^ resulting in ROS-induced oxidative damage that drives genotoxicity ^12^ and DNA breaks ^13^. Current prevention strategies for EAC development in patients with Barrett’s esophagus, such as chemoprevention with proton pump inhibitors, aspirin, and statins, as well as endoscopic surveillance, show only modest and inconsistent effectiveness^14^. This highlights the need for the development of novel therapeutic strategies to counteract the carcinogenic effects of acid-biliary reflux.

The endocannabinoid system (ECS) is a complex network of lipid-based neurotransmitters, receptors (primarily CB1 and CB2), and enzymes that regulate their synthesis and degradation, playing a central role in maintaining homeostasis across multiple organ systems, including the gastrointestinal, immune, metabolic, and cardiovascular systems ^15^. Exogenous phytochemicals from *Cannabis sativa* can mimic or modulate many of the same physiological effects as produced by endogenous endocannabinoids such as anandamide (AEA) by interacting with the same cannabinoid receptors within the ECS ^16^. Cannabinoids and terpenes, the two principal classes of phytochemicals derived from *Cannabis sativa*, both exhibit anti-inflammatory properties and have been explored therapeutically for chronic disorders such as multiple sclerosis and irritable bowel syndrome ^17-19^. Cannabinoids such as cannabigerol (CBG) can modulate intracellular levels of ROS and the natural antioxidant superoxide dismutase ^15,20^. Terpenes, aromatic compounds also abundant in *Cannabis sativa*, target inflammatory signaling pathways and display moderate anti-inflammatory effects, in part by modulating the levels of TNFα and IL-1β ^21^. *In vitro* evidence suggests that phyto-cannabinoids and terpenoids inhibit proliferation in multiple cancer cells^22,23^. Notably, whole-plant cannabis extracts, which combine cannabinoids and terpenes, often show greater efficacy than isolated compounds, supporting the concept of a synergistic “entourage effect” ^24^. The entourage effect describes the potential for cannabinoids, terpenes, and other cannabis-derived compounds to work synergistically, possibly enhancing therapeutic effects compared to isolated cannabinoids; however, the optimal ratio combinations and the specific diseases or conditions best targeted by these combinations remain unknown ^25-27^

Here, we systematically evaluated various cannabinoid-terpene combination ratios to elucidate their synergistic antioxidant effects in physiologically relevant models of GERD. Additionally, we explored whether specific cannabinoid-terpene ratios could mitigate DCA-induced mutagenesis and DNA damage through an entourage effect. These preliminary findings support future *in vivo* investigation and provide proof of concept that targeting the ECS with phytochemicals may offer novel therapeutic strategies for the prevention of EAC.

## Material and Methods

### Chemicals and reagents

Cannabinoids were purchased from Cayman Chemical and terpenes were purchased from Sigma Aldrich. Bile acid cocktail consisted of an equimolar mixture of glycocholate, taurocholate, glycodeoxycholate, glycochenodeoxycholate, and deoxycholate at a final concentration of 300 µM. This cocktail reflects the mixture of bile acids to which the distal esophagus is ordinarily exposed during gastroesophageal reflux ^28^.

### Cell Culture

Human cell lines were purchased from the following vendors: HET1a (ATCC, cat# CRL-2692, Virginia, USA), Human Esophageal Epithelial Cells (Sciencell, cat# 2720, California, USA; CellBiologics, cat# H-6046, Illinois, USA), CP-A (KR-42421) (ATCC, cat # CRL-4027, Virginia, USA). According to the vendor information, the cells were obtained under the IRB-approved protocols. Normal esophageal cells were plated on flasks pre-coated with 0.01 mg/mL human fibronectin (Corning, cat# 356008, New York, USA) and 0.03 mg/mL Collagen I, bovine (ChemCruz, cat# sc-29009, Texas, USA) and grown in Bronchial Epithelial Cell Growth Medium (Lonza, Cat# CC-3170, Basel, Switzerland). CP-A Barrett’s esophagus cells were maintained in MCDB-153 supplemented with 0.4 µg/ml hydrocortisone, 20 ng/ml recombinant human epidermal growth factor, 8.4 µg/L cholera toxin, 20 mg/L adenine, 140 µg/ml bovine pituitary extract, 1x ITS Supplement (Sigma; I1884), 4 mM glutamine, and 5% fetal bovine serum.

### GPCR gene expression analyses

These analyses were performed using publicly available data in NCBI Gene Expression Omnibus under GSE1420^29^. No active patient enrollment requiring an IRB approval was performed in this study. The gene expression data from the Affymetrix Human Genome U133A Arrays, comprised of the normal esophageal epithelium (n=8), Barrett’s esophagus (BE) (n=8), and esophageal adenocarcinomas (EAC) (n=8), were analyzed using the GEO2R interactive web tool.

### Cell viability analysis

Cells were cultured prior to exposure to the indicated test articles. After treatment, cells were washed and resuspended in a phenol red-free RPMI or DMEM and subsequently treated with the XTT assay (ThermoFisher, Waltham, MA, USA) or the MTS assay (Promega, Madison, WI, USA) following manufacturer protocols.

### Immunohistochemical Staining

To assess the expression of CB1, a tissue array containing 50 cases/50 cores of esophageal adenocarcinoma, cardia adenocarcinoma, and normal esophageal and cardia tissue was obtained from Biomax (cat# BC001113, Rockville, MD, USA). The tissue sections were then rehydrated through a series of ethanol solutions and placed in distilled water. Antigen retrieval was performed in sodium citrate, pH 6.0 (Sigma, S-4641, St. Louis, MO, USA) using an electric pressure cooker. The tissue sections were placed in a solution of 0.1% TritonX-100 (Sigma, T9284) in phosphate-buffered saline (PBS) for 15 minutes and pre-blocked with hydrogen peroxide Blocking Reagent (Abcam, 64218, Cambridge, UK). Blocking solution consisting of 10% normal donkey serum in PBS (EMD Millipore, S30-100 ML, Billerica, MA, USA) was added to the slides and left for 30 minutes at room temperature. The blocking solution was replaced with the primary antibodies diluted in blocking solution using rabbit anti-CB1 (clone D5N5C, Cell Signaling Technologies, cat# 93815, Massachusetts, USA) and anti-epithelial cell adhesion molecule (EPCAM) (Origene, cat# UM500096, Maryland, USA), which was used to mark EAC. The tissue sections were incubated in primary antibody solution overnight at 4°C, and washed twice with 0.1% Tween-20 (Promega, H5151, Madison, WI, USA) in PBS. Sections were then incubated with secondary antibody (donkey anti-rabbit IgG-594 and donkey anti-mouse IgG-488) diluted 1:500 in PBS for 1 hour at RT and washed twice in 0.1% Tween-20 in PBS for 15 minutes each. 4′,6-diamidino-2-phenylindole (DAPI) was used to stain nuclei. Semiquantitative analysis was performed using an H-score analysis by two independent observers. The proportion (0-100) and intensity of CB1 immunostaining (0: no staining; 1: weak staining; 2: moderate staining, 3: strong staining) were used to calculate an H-score.

### Mitochondrial Membrane Potential

MitoProbe JC-1 Assay Kit for Flow Cytometry (cat# M34152, ThermoFisher, Massachusetts, USA) was used to measure the mitochondrial membrane potential. Depending on the experimental group, cells were pre-treated with the CBG/Phytol admixture or DMSO and incubated for 2 hours at 37°C in a CO_2_ incubator. Cells were then treated with DCA with the CBG/Phytol admixture or DMSO. After treatment, cells were loaded with 2µM of JC-1 and incubated for 15 minutes at 37°C. Cytoplasmic JC-1 monomers were detected in the green spectrum (∼529 nm) while mitochondrial J-aggregates were detected in the red spectrum (∼590 nm). Results are representative of 2 and 3 independent repeats per cell line.

### Detection of intracellular reactive oxygen species (ROS)

Prior to treatments, cells were washed and incubated with 2 μM CM-DCFDA (Life technologies, Grand Island NY) for 10□minutes followed by a wash in PBS and then recovery in DMEM for 15□min. Cells were then treated as described in the figure legend. After treatment, cells were analyzed by fluorescent plate reader or trypsinized to single cells and processed by flow cytometry (excitation: 488 nm; emission: 535 nm). Fluorescence intensity was calculated as % increase of vehicle control.

### DNA Damage

The activation of ATM and H2AX (a marker of DNA breaks) was measured using the Muse Multi-Color DNA Damage kit (MilliporeSigma, Burlington, MA, USA) according to the manufacturer’s instructions. The percentage of ATM-activated cells and H2AX-activated cells was determined as dual activation by monitoring expression of both the ATM and γ-H2AX, using the Muse Cell Analyzer (MilliporeSigma, Burlington, MA, USA). Expression of γ-H2AX was determined by flow cytometry on an Accuri C6 flow cytometer following manufacturer protocol (MilliporeSigma, Burlington, MA, USA).

### Evaluation of impact on genome stability

HET1A cells were cultured with either vehicle, 100µM DCA, CBG/Phy admixture, or a combination of DCA and CBG/Phy admixture for 14 days. DNA from these and parental (Day 0) cells was extracted using a QIAGEN DNeasy Blood & Tissue Kit (Qiagen, cat# 69504, Maryland, USA) and hybridized to PMDA arrays (Affymetrix). Genomic instability in cultured cells was assessed by identifying new copy number events (both deletions and amplifications), using the DNA of “Day 0” cells as a reference point.

## Results

### CBG and Phytol (Phy) at a 1:5 ratio counteract DCA-mediated mitochondrial depolarization and DNA damage in normal esophageal epithelial cells

To explore whether cannabinoids and terpenes can minimize the harmful effects of DCA, we conducted cell viability, mitochondrial depolarization, and DNA damage assays using an established esophageal epithelial cell line, HET1a,^30^ and primary human esophageal epithelial cells HEsEpiC as summarized in **Fig. 1A**. Using various DCA concentrations previously found in the esophageal aspirates of GERD patients^31^, we determined the lethal dose 50% (EC50) for DCA in HET1a cells to be 295.4 µM with an R^2^ of 0.9935 (**Fig. S1A**). We also found that DCA can induce mitochondrial membrane depolarization at doses equal or higher than 300 µM (**Fig. 1B**). We tested combinations of cannabinoids and terpenes with known antioxidative properties for their ability to counteract the DCA-induced DNA damage (**Fig. 1C**). In HEsEpiC cells treated with DCA, we observed more than 5% increase in ATM+ H2AX+ cells compared to untreated controls (**Fig. 1C**). The addition of the tetrahydrocannabinoid (THC) had no effect on the DCA-induced DNA damage, while the combination of CBG and the terpenoid Phy resulted in a significant reduction in the percentage ATM+ H2AX+ cells (**Fig. 1C and Fig. S2**). Of note, similar effect was observed when cells were treated with CBG in combination with myrcene or β-caryophyllene (**Fig. 1C and Fig. S2)**. Additionally, we interrogated various CBG and Phy concentrations for their ability to reduce the DCA-induced ROS (**Fig. S1A**). We found that the combination of 1 µM CBG and 1 µM Phy was capable of significantly decreasing ROS compared to the untreated control (p < 0.0001). Subsequently, we tested several CBG and Phy combination ratios for their ability to reverse DCA-induced mitochondrial membrane depolarization and DNA damage. We determined that the 1:5 combination ratio of CBG and Phy can reverse DCA-induced mitochondrial membrane depolarization (**Fig. 1D**). In addition, the 1:5 CBG/Phy combination ratio had a trend to reduce the DCA-triggered formation of DNA breaks in the HET1a cell line, measured as a percentage of ATM+H2AX+ cells (**Fig. 1E**).

**Figure 1.**
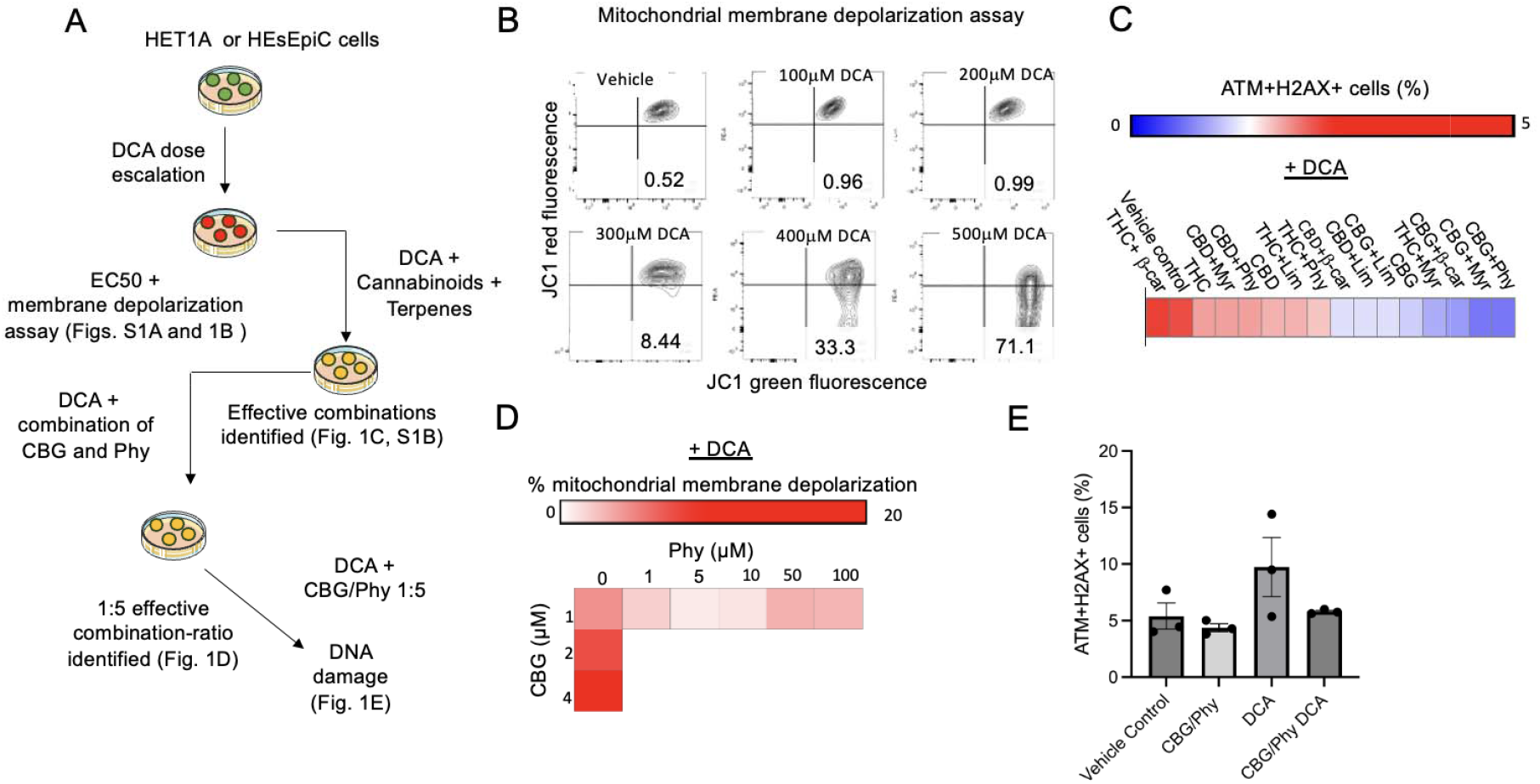
A CBG and Phy combination ratio of 1:5 can reverse DCA-induced mitochondrial depolarization and DNA damage in HET1a and HEsEpiC esophageal epithelial cells. (A)Schematic illustration of experimental design. (B)Representative flow cytometry analyses of mitochondrial membrane potential in DCA-treated HET1A cells measured by red- and green-fluorescent changes performed using the MitoProbe JC-1 Assay Kit. Cells with normal mitochondrial membrane potential are depicted in the right upper corner (red), and the cells with reduced mitochondrial membrane potential are shown in the right lower corner (green). (C)The heatmap illustrates the percentage of ATM+/H2AX+ cells in HEsEpiC cell cultures pretreated with diverse cannabinoids and terpenes at a 1:5 ratio, while exposed to 100 µM DCA. Data were analyzed using the Muse Multi-color DNA Damage Kit (Cytek, USA). (D)The heatmap illustrates the changes in the mitochondrial membrane potential in HET1A cells treated with varied concentrations and ratios of CBG and Phy in the presence of 100 µM DCA. (E)The bar graph illustrates the percentage of ATM+/H2AX+ HET1A cells after treatment with CBG and Phytol at 1:5 ratio in the presence of DCA. Data were analyzed using the Muse Multi-color DNA Damage Kit (Cytek, USA).

### CBG and Phy admixture restricts proliferation and promotes apoptosis of DCA-damaged esophageal cells

To test whether CBG/Phy can prevent propagation of the DCA-damaged esophageal epithelial cells, we first assessed their effect on cell proliferation following DCA exposure. HET1A cells were pretreated with CBG/Phy at a 1:5 ratio and exposed to DCA for 24 hours. Subsequently, the cells were incubated with fresh media without DCA and with CBG/Phy or vehicle control. At 48 hours of culture, the XTT cell viability assay revealed that CBG/Phy treatment alone had no significant effect on cell proliferation compared to untreated controls. In contrast, DCA significantly inhibited HET1A cell proliferation at 48 hours, however, a significant increase in cell proliferation was subsequently observed at 72 hours of culture, i.e. 48 hours after DCA removal **(Fig. 2A)**. In contrast, HET1A cultures pre-treated with CBG/Phytol admixture and exposed to DCA exhibited reduced proliferation even after DCA withdrawal (**Fig. 2A**). These results indicate that while the cells treated with DCA have the ability to resume cell proliferation after DCA removal and, thus propagate cells with acquired DNA alterations, treatment with CBG/Phy admixture can prevent expansion of these damaged cell populations. To test whether CBG/Phy treatment sensitizes DCA-damaged cells to apoptosis, we exposed HET1A cells to DCA concentrations ranging from 0 to 500 µM, with CBG/Phy at a 1:5 ratio or vehicle control pre-treatment (**Fig. 2B**). We found that CBG/Phy reduced DCA EC50 to 281.3 µM from 370.6 µM observed in the vehicle control group. Taken together, these results suggest that the CBG/Phy combination prevents the proliferation and induces apoptosis of DCA-damaged cells, which could be critical for disrupting esophageal carcinogenesis in the setting of DCA exposure.

**Figure 2.**
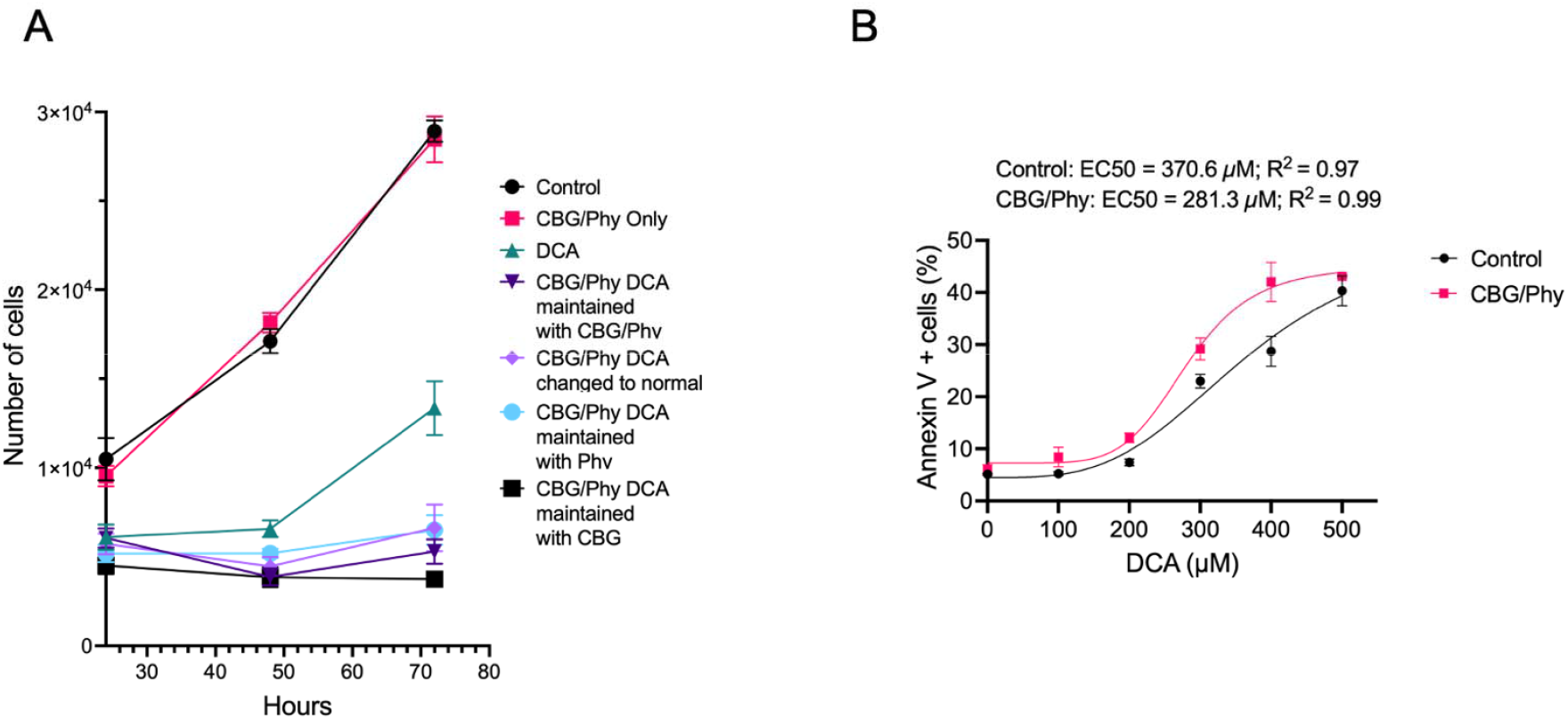
The effect of CBG and Phy on the proliferation and apoptosis of DCA-damaged esophageal cells. (A)XTT cell viability assay of HET1a cells subjected to 100 µM DCA or vehicle control treatment for 24 hours in the presence or absence of CBG and P. (B)The plot represents a non-linear regression analysis of the percentage of apoptotic cells in the HET1a cell line cultured with or without CBG/Phy pre-treatment and exposed to DCA at concentrations ranging from 0 to 500 µM.

### CBG and Phy Attenuate Genomic Instability Triggered by DCA

Based on the previously reported mutagenic effects of DCA ^32^ and our current results demonstrating the role of the CBG/Phy 1:5 admixture in counteracting DCA-induced DNA damage, we assessed the role of CBG/Phy in reversing genomic instability caused by DCA using the whole genome PMDA arrays (Affymetrix) (**Fig. 3A**). We found that treatment of HET1A cells with DCA resulted in a significant acquisition of new copy number events over a period of three weeks compared to the vehicle control (**Fig. 3A**). Specifically, we observed an increase in both gene amplifications and deletions (**Fig. 3B**). When HET1A cells were treated with the 1:5 CBG/Phy combination after DCA exposure, we observed a significant decrease in gene amplifications and deletions (**Fig. 3B**). Notably, no significant changes in the new copy number events were observed with the CBG/Phy treatment alone in the absence of DCA compared to the vehicle control samples (**Fig. 3B**). These data demonstrate that treatment with CBG/Phy can attenuate DCA-triggered genomic instability and reduce mutagenicity in the setting of chronic DCA exposure.

**Figure 3.**
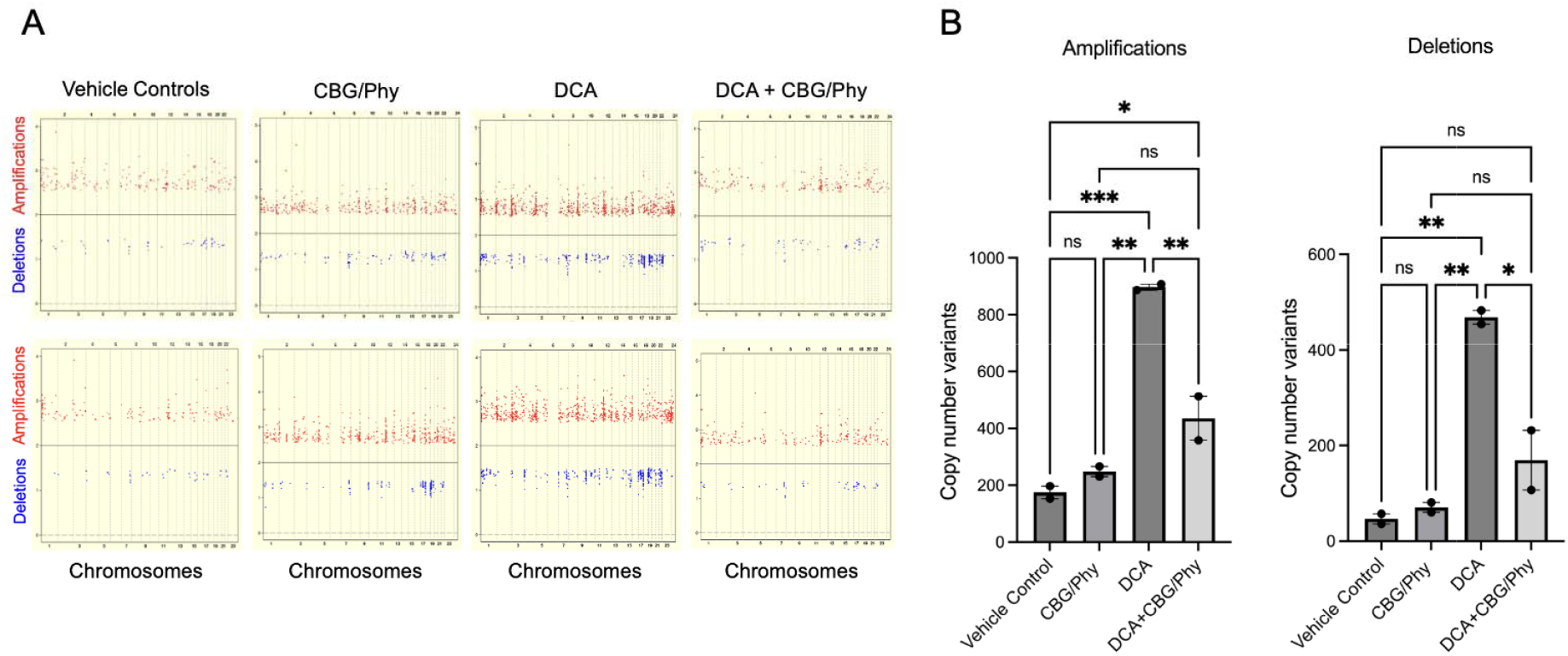
CBG and Phy at a 1:5 ratio reduce DCA-induced genomic instability in normal esophageal epithelial cells. (A)Scatter plots of the whole genome amplifications and deletions in HET1A cells cultured in the presence of either vehicle control, 100 µM DCA, CBG/Phytol admixture, or a combination of 100 µM DCA and CBG/Phytol admixture for 14 days. DNA extracted at day 14 was compared using PMDA arrays (Affymetrix) to the DNA of untreated cells collected at day 0. (B)Bar graphs illustrate the quantitative analyses of the whole genome amplifications (left panel) and deletions (right panel). Data were analyzed using one-way ANOVA with multiple comparisons. * p < 0.05,**p < 0.01, ***p < 0.001; ns, not significant.

### CBG/β-caryophyllene 1:5 combination mitigates low pH and bile acid-induced ROS in metaplastic Barrett’s esophagus cells

Next, we asked whether the combinations of cannabinoids and terpenes can also protect metaplastic esophageal cells following acute exposure to a low pH environment (pH 4.5) combined with a bile acid cocktail comprised of multiple secondary bile acids at concentrations mimicking the caustic environment during gastroesophageal reflux (GER) ^31^ (**Fig. 4A**). The Barrett’s esophagus cell line, CP-A, was pre-treated with either a vehicle control or cannabinoids in the presence or absence of terpenes for 12 hours, followed by an acute (15 minutes) exposure to low pH and bile acid cocktail (**Fig. 4A**). The cells were either immediately analyzed for ROS via fluorescent detection of CM-DCFDA or recovered for 24 hours and analyzed for DNA damage via γ-H2AX flow cytometry. We found that acute exposure to the low pH/bile acid combination results in significant increase in ROS **(Fig. 4B)**. Using an MTS cell viability assay, we determined that 1 µM cannabinoid did not result in increased cell death or affected proliferation (**Fig. 4C**).

**Figure 4.**
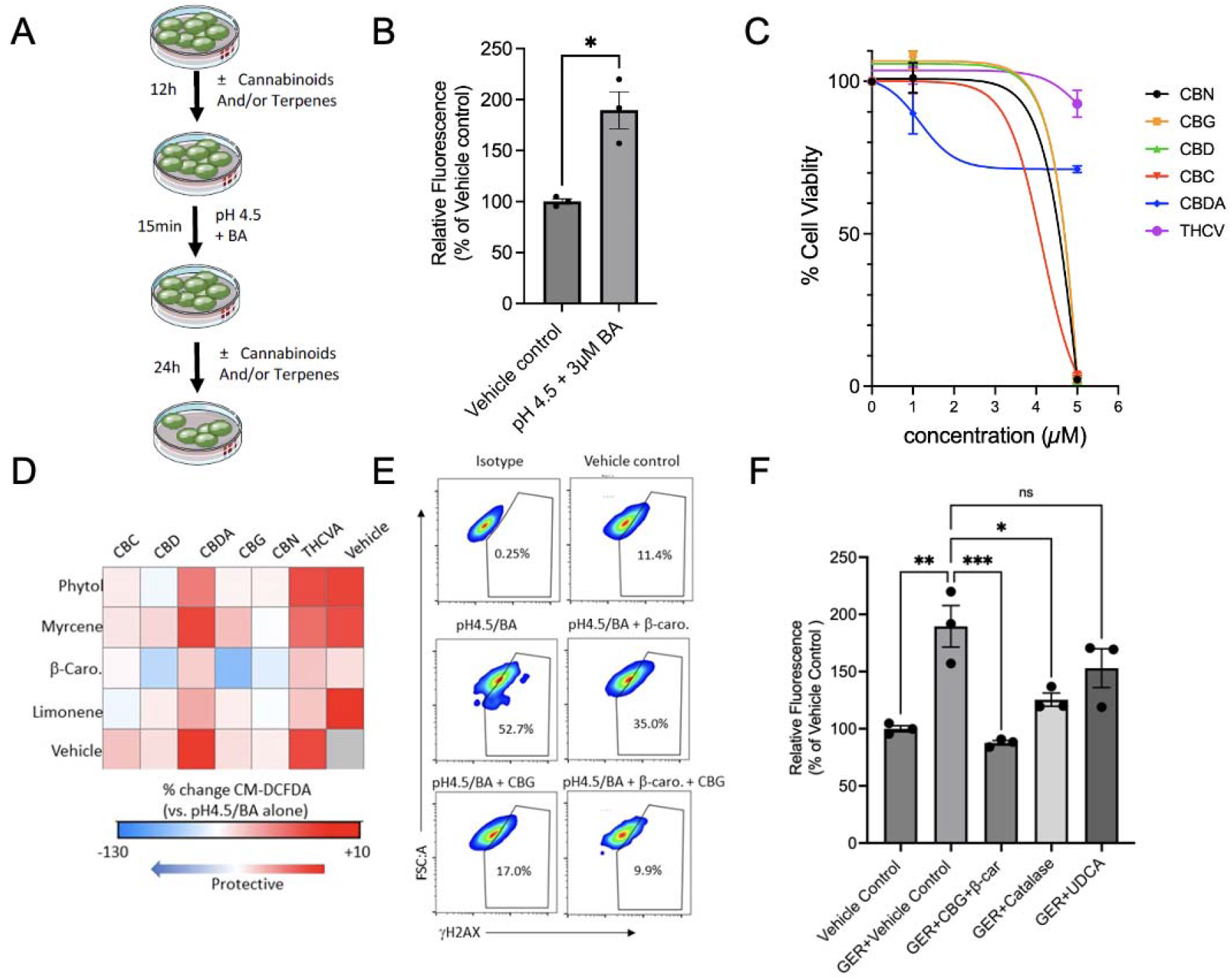
CBG and β-car (1:5) reverse the damaging effect of the acid-containing gastroesophageal refluxate (GER) in metaplastic Barrett’s esophageal cells. (A)Experimental design schematic. BE cell line CP-A was treated with a vehicle control or cannabinoids in the presence or absence of terpenes for 12 hours, followed by an acute 15-minute exposure to a physiologically relevant low pH and bile acid cocktail, as described in the Methods. The cells were either immediately analyzed for ROS via fluorescent detection of CM-DCFDA or recovered for 24 hours and analyzed for DNA damage using γ-H2AX flow cytometry. (B)Bar graph illustrates the comparative analyses of ROS measured by CM-DCFDA fluorescence in CP-A cells following acute exposure to GER. The samples were analyzed by flow cytometry, and the resu ts are shown as a percent increase compared to the vehicle control. The data were analyzed using an unpaired t-test with Welch’s correction. * *p* < 0.05, n=3 (C)Non-linear fit of the dose response cell viability analysis of CP-A cells after exposure to cannabinoids for 24 hours, analyzed using the MTS Assay Kit. (D)Heat map shows changes in ROS induced by GER in CP-A cells pre-treated with the cannabinoid and terpene combinations at 1:5 ratio as determined by CM-DCFDA fluorescence. Data are expressed as the percent change when compared to GER alone. (E)Representative flow cytometry analyses of DNA damage as determined by γ-H2AX fluorescence of CP-A cells pre-treated with CBG and β-car following exposure to GER. (F)Bar graph depicts the comparative analyses of ROS measured by CM-DCFDA fluorescence in CP-A cells exposed to GER and antioxidants (Catalase), ursodeoxycholic acid (UDCA), or the cannabinoid/terpene combination of CBG and β-car at a 1:5 ratio. The samples were analyzed by flow cytometry, and the results are shown as a percent increase compared to the vehicle control. The data were analyzed using one-way ANOVA with Šídák’s multiple comparisons test. * *p* < 0.05, \*\**p* < 0.01, \*\*\**p* < 0.001; ns, not significant.

Next, using flow cytometry to screen various combinations of terpenes and cannabinoids at the 1:5 ratio, we determined that, while CBG and Phy reduced the amount of ROS caused by the low pH/bile acid insult, the highest reduction in ROS was identified in the combination of CBG and β-caryophyllene (β-car) **(Fig. 4D)**. Indeed, when we tested this combination in the residual cells, harvested 24 hours post-treatment, we determined a significant diminishment of DNA damage, as determined by γ-H2AX fluorescence **(Fig. 4E)**. Lastly, we compared other ‘gold standard’ combinations that are known to reduce the toxic assault of GER including catalase and the tertiary bile acid ursodeoxycholic acid (UDCA)^33^. Notably, the results suggested that CBG/β-car (1:5) treatment mitigated the oxidative insult of GER better than UDCA (**Fig. 4F**).

### Association of CB1 and G-protein coupled receptors (GPCRs) with esophageal carcinogenesis

It has become increasingly clear that the ECS functions through multiple receptor pathways in addition to CB-1 and CB-2 ^34^. Indeed, some GPCRs, such as GPR35^35^ and GPR63^36^, are thought to have affinity for endo- and phyto-cannabinoids ^35^. To evaluate the role of GPCRs in the BE-associated carcinogenesis, we first examined the expression of the established cannabinoid G protein-coupled receptor CB-1 ^37^ in normal esophagus and EAC. We observed punctate CB1-expression mainly in the basal layer of the normal esophageal stratified squamous epithelium **(Fig. 5 A&B)**. In EAC, CB1 was co-expressed with the epithelial marker EPCAM (**Fig. 5 C&D**) and was significantly upregulated based on the H-score semiquantitative intensity analysis (**Fig. 5E**). Next, we investigated the expression of additional GPCRs in normal epithelium, BE metaplasia, and esophageal adenocarcinoma (EAC) from the publicly available data GSE1420^29^. Using the GEO2R interactive web tool, based on the log2FC > 0.9 and adjusted p-values of <0.05, we identified a cohort of GPCRs that were significantly differentially expressed in BE and EAC compared to normal esophageal epithelium. Among them, GPR35 and GPR5CA were significantly upregulated in both BE (**Fig. 5A**) and EAC (**Fig. 5B**), while GPR63 was downregulated in BE (**Fig. 5A**). GPR45 and GPR143 were downregulated specifically in EAC (**Fig. 5B**). These observations point to the potential role of modulating CB1 and additional GPCRs for the prevention and treatment of EAC.

**Figure 5.**
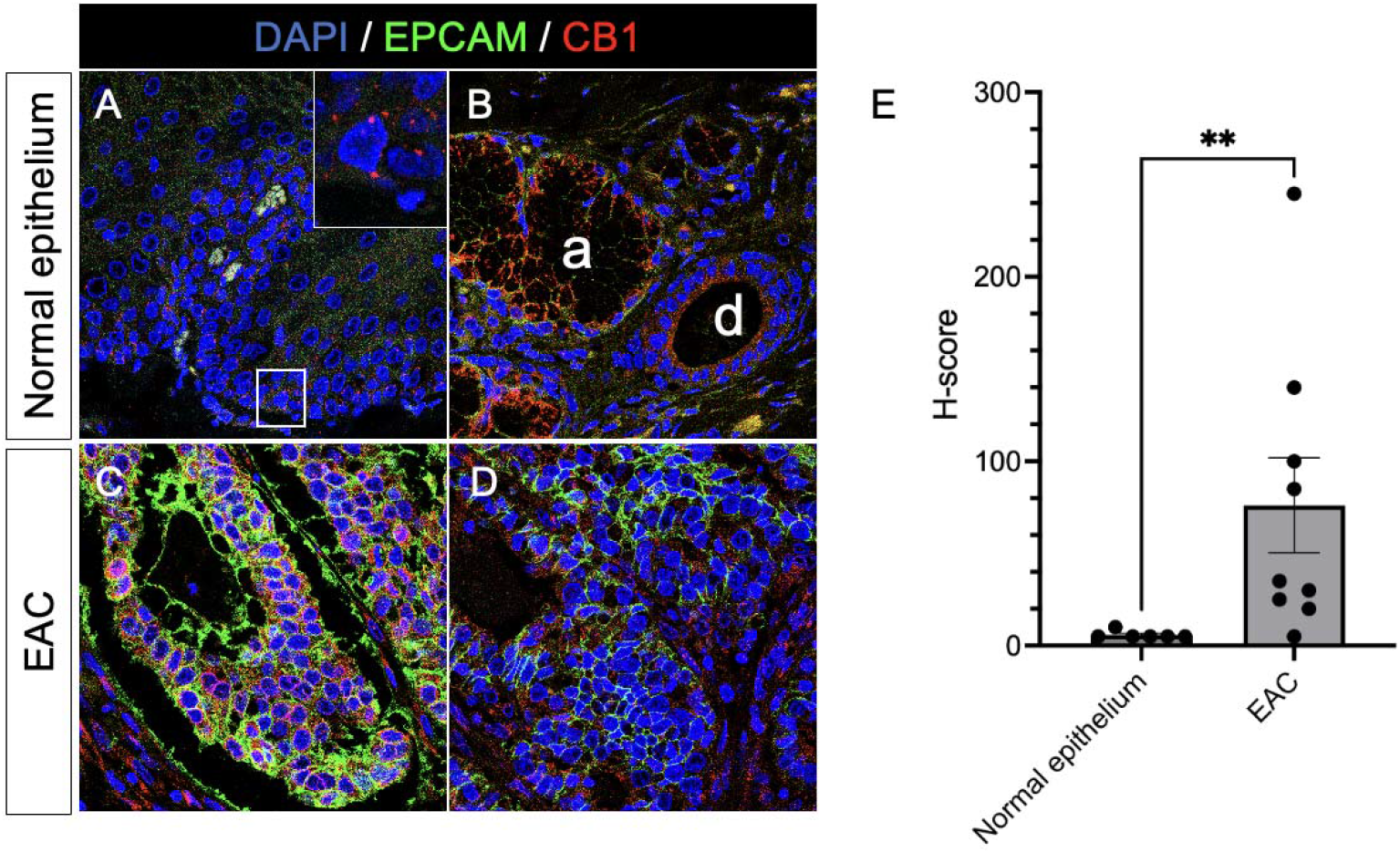
Expression of CB1 in human normal esophageal mucosa and EAC assessed by confocal microscopy. (A)A representative immunofluorescence analysis of CB1 (red) and EPCAM (green) expression in the normal human esophageal mucosa (Magnification 63x). The inset highlights the CB1 expression in the basal epithelial layer. (B)A representative immunofluorescence analysis depicting CB1 and EPCAM co-expression in the ductal epithelium of the submucosal gland (d) and acini of the submucosal gland (a). (C&D)Representative immunofluorescence analyses of CB1 and EPCAM co-expression in two distinct human EAC samples. **(E)**Bar graph represents a comparative H-score semiquantitative intensity analysis of CB1 expression in human normal esophageal mucosa and EAC. The data were analyzed using the Mann-Whitney U test. ** *p* < 0.01.

**Figure 6.**
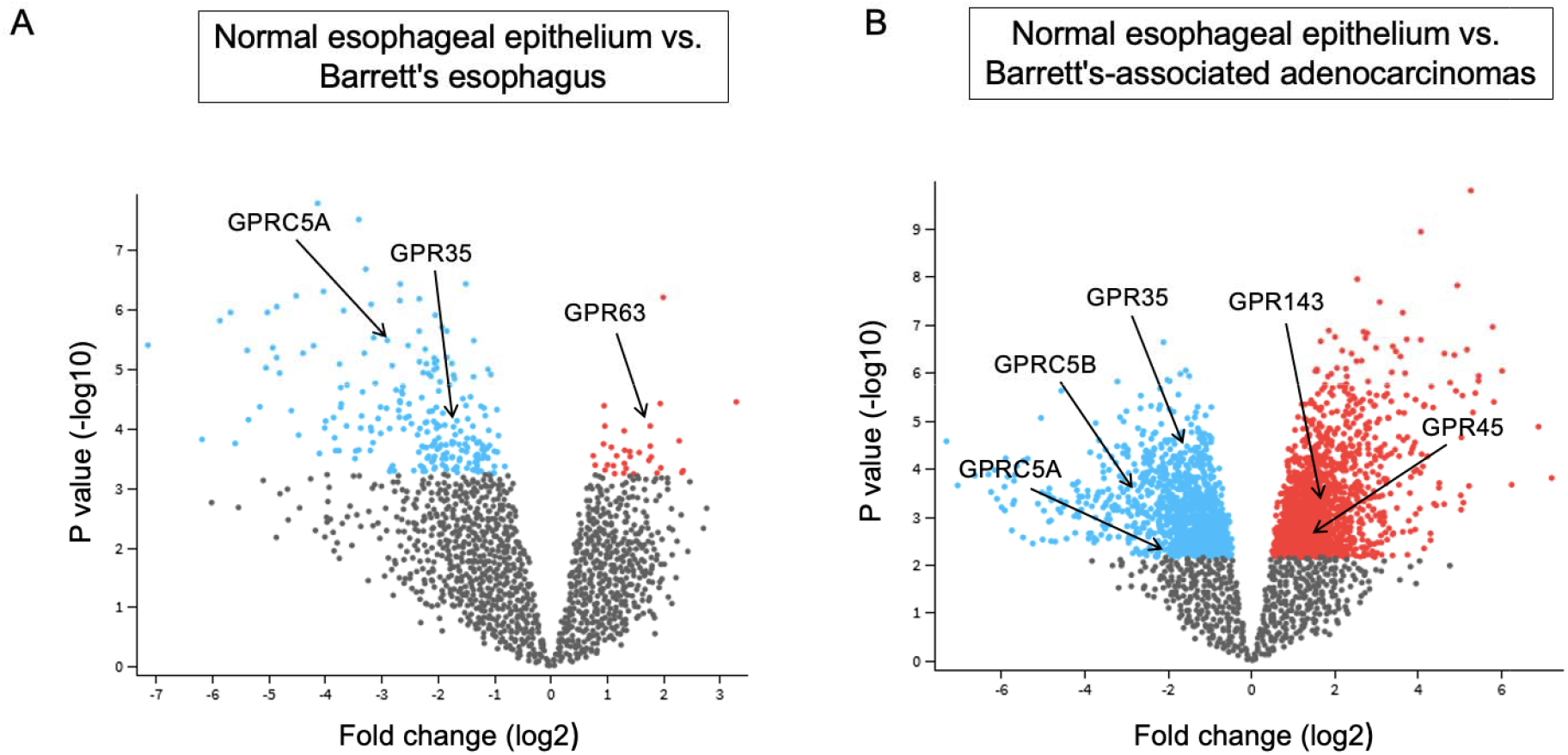
Dysregulation of GPCR gene expression in clinical BE and BE-associated EAC. (A)The volcano represents differentially expressed genes in clinical BE (n=8) compared to the normal esophageal epithelium (n=8). The data were obtained from the GSE database under GSE1420. BE upregulated (blue) and downregulated (red) genes were identified using the GEO2R tool, based on log2FC> 0.9 and adjusted p-values of <0.05. (B)The volcano represents differentially expressed genes in clinical BE-associated EAC (n=8) compared to the normal esophageal epithelium (n=8). The data were obtained from the GSE database under GSE1420. EAC upregulated (blue) and downregulated (red) genes were identified using the GEO2R tool, based on log2FC > 0.9 and adjusted p-values of <0.05

## Discussion

We present a first-of-its-kind drug screening effort for GERD-induced damage that leverages the properties of the ECS using bioactive phyto-cannabinoids and terpenes. We discovered that specific combinations and ratios of these combinations are effective for different pathologic etiologies in the esophagus. It remains to be studied how these combinations and ratios can be deployed in a clinical manner to prevent the conversion of malignant disease and suppress the mutagenic effects of bile acid refluxate.

Leveraging the medicinal properties of natural products is not a new concept. There are many bioactive compounds present in medicinal plants that have shown potential for cancer therapy. In particular, bioactive compounds, such as [6]-Gingerol, thymoquinone, artepillin C, and *Gaboderma atrum* polysaccharide, which have been shown to disrupt the mitochondrial membrane potential, induce release of cytochrome C, and activate caspase activity ^38^. On the other hand, the plant *Cannabis sativa* has been used as a ‘medicinal’ plant for thousands of years ^39^, yet the medicinal properties have been poorly studied, given the governmental and legal issues surrounding it, globally ^40,41^. In contrast to other natural products and their individual activity, we found that cannabinoids and terpenes share synergistic qualities that require specific combinations in order to be ‘therapeutic’, at least in the context of GER and GERD. There are many more combinations that could be tested, including three-compound and four-compound combinations, among others.

Our findings support the hypothesis that CBG and Phy counteract the DCA-mediated increase of ROS production and ensuing oxidative DNA damage. Therefore, we investigated the potential to use the CBG/Phy admixture as a chemopreventive therapy. However, our findings also show that the CBG/Phy admixture primes human esophageal epithelial cells to undergo apoptosis when exposed to high concentrations of DCA. Our data showed that in human esophageal cells, pre-treatment with the CBG/Phytol admixture and 300 µM DCA led to a significant disruption of the mitochondrial membrane potential and increased cellular apoptosis. This may explain why preincubation with CBG/Phytol was required to counteract the effect of DCA and why it could significantly counteract DCA-mediated genomic instability. Our findings that CBG and β-car promote an optimal therapeutic response in metaplastic cells following low pH and bile acid insult suggest that one combination is not universally protective and may even be disease-specific.

*Proposed mechanism of action*. We uncovered that the CBG and either Phy or β-car admixture affected the long-term recovery potential of human esophageal epithelial cells. We showed that 100 µM of DCA is sufficient to affect cell proliferation but does not induce mitochondrial-mediated cell apoptosis. Mechanistically, it has been suggested that unconjugated bile acids disrupt cellular pathways and promote the release of ROS from the mitochondria by perturbing the mitochondrial outer membrane and promoting mitochondrial swelling ^42^. CBG can activate mitochondrial CB1 through intramitochondrial G-alpha-i and inhibit soluble adenylyl cyclase, which inhibits protein kinase A and leads to downstream phosphorylation of proteins involved in the mitochondrial electron transport system ^43^. Phy, for example, regulates cellular respiration by irreversibly inhibiting SSADH, which is present in the inner mitochondrial membrane ^44^. Our data show that maintaining cells with the CBG/terpene and DCA showed reduced cellular proliferation, which could be the result of long-term damage to the mitochondria. Moreover, the CBG/terpene admixture can potentially inhibit/reverse DCA-induced DNA instability by promoting mitochondrial membrane instability and subsequent mitochondrial-mediated apoptosis. Therefore, the CBG/terpene admixture may promote a reduction in total DNA damage/instability by inducing apoptosis and eliminating cells with significant DNA instability. A potential upside of clearing DCA-mediated damaged mitochondria is that it may help counteract DCA-mediated genomic instability. There is a possibility that another mechanism is responsible for the effect on cellular viability by the admixture, but it will require further investigation. It is important to note that additional evaluation in models that more closely represent human physiology, including multi-cell complex in-vitro models and *in-vivo* models, should be evaluated to confirm the effect of our proposed mechanism of action and efficacy for combinations of CBG and terpenes.

Together, the data suggest that hereby identified CBG and either Phy or β-car admixtures promote mitochondrial-mediated cell apoptosis only in combination with DCA. This further suggests that these admixtures prime cells for apoptosis and furthermore interfere with a mutation-riddled recovery. A combination of CBG and terpenes may lead to novel cancer prevention strategies to overcome the significant rise in esophageal cancer, a disease for which no effective treatment exists.

## Author contributions

A.G. and G.G. designed and performed in-vitro experiments and analyzed data. S.A.K., M.A.S., and L.B. assisted with experiments. H.M., M.H.F., and N.Y.F. supervised the project. A.G., G.G., and N.Y.F. wrote and revised the paper. All authors reviewed and approved the final manuscript.

## Data sharing statement

The datasets generated during and/or analyzed in the current study are available from the corresponding author upon reasonable request.

## References

1. Falk, G. W. et al. Barrett’s esophagus: prevalence-incidence and etiology-origins. Ann. N. Y. Acad. Sci. 1232, 1–17, doi:10.1111/j.1749-6632.2011.06042.x (2011).

2. Shields, H. M. et al. Barrett’s esophagus: prevalence and incidence of adenocarcinomas. Ann. N. Y. Acad. Sci. 1232, 230–247, doi:10.1111/j.1749-6632.2011.06054.x (2011).

3. Saha, B. et al. Prevalence of Barrett’s Esophagus and Esophageal Adenocarcinoma With and Without Gastroesophageal Reflux: A Systematic Review and Meta-analysis. Clin Gastroenterol Hepatol 22, 1381–1394 e1387, doi:10.1016/j.cgh.2023.10.006 (2024).

4. Wang, S., Li, Z., Zhou, Z. & Kang, M. Causal analysis of gastroesophageal reflux disease and esophageal cancer. Medicine (Baltimore) 103, e37433, doi:10.1097/MD.0000000000037433 (2024).

5. Quante, M., Graham, T. A. & Jansen, M. Insights Into the Pathophysiology of Esophageal Adenocarcinoma. Gastroenterology 154, 406–420, doi:10.1053/j.gastro.2017.09.046 (2018).

6. Fujimura, T. et al. Inflammation-related carcinogenesis and prevention in esophageal adenocarcinoma using rat duodenoesophageal reflux models. Cancers (Basel) 3, 3206–3224, doi:10.3390/cancers3033206 (2011).

7. Flejou, J. F. Barrett’s oesophagus: from metaplasia to dysplasia and cancer. Gut 54 Suppl 1, i6–12, doi:10.1136/gut.2004.041525 (2005).

8. Bernstein, H., Bernstein, C., Payne, C. M., Dvorakova, K. & Garewal, H. Bile acids as carcinogens in human gastrointestinal cancers. Mutat. Res. 589, 47–65, doi:10.1016/j.mrrev.2004.08.001 (2005).

9. Rodrigues, C. M., Fan, G., Ma, X., Kren, B. T. & Steer, C. J. A novel role for ursodeoxycholic acid in inhibiting apoptosis by modulating mitochondrial membrane perturbation. J. Clin. Invest. 101, 2790–2799, doi:10.1172/JCI1325 (1998).

10. Sousa, T. et al. Deoxycholic acid modulates cell death signaling through changes in mitochondrial membrane properties. J. Lipid Res. 56, 2158–2171, doi:10.1194/jlr.M062653 (2015).

11. Goldman, A. et al. The Na+/H+ exchanger controls deoxycholic acid-induced apoptosis by a H+-activated, Na+-dependent ionic shift in esophageal cells. PLoS One 6, e23835, doi:10.1371/journal.pone.0023835 (2011).

12. Cadenas, E. & Davies, K. J. Mitochondrial free radical generation, oxidative stress, and aging. Free Radic. Biol. Med. 29, 222–230, doi:10.1016/s0891-5849(00)00317-8 (2000).

13. Karanjawala, Z. E., Murphy, N., Hinton, D. R., Hsieh, C. L. & Lieber, M. R. Oxygen metabolism causes chromosome breaks and is associated with the neuronal apoptosis observed in DNA double-strand break repair mutants. Curr. Biol. 12, 397–402, doi:10.1016/s0960-9822(02)00684-x (2002).

14. Abrams, J. A. Are We Making Progress in Preventing Barrett’s-Related Esophageal Cancer? Therap Adv Gastroenterol 2, 73–77, doi:10.1177/1756283X08102044 (2009).

15. Borrelli, F. et al. Beneficial effect of the non-psychotropic plant cannabinoid cannabigerol on experimental inflammatory bowel disease. Biochem. Pharmacol. 85, 1306–1316, doi:10.1016/j.bcp.2013.01.017 (2013).

16. Fisar, Z. Phytocannabinoids and endocannabinoids. Curr Drug Abuse Rev 2, 51–75, doi:10.2174/1874473710902010051 (2009).

17. Borgelt, L. M., Franson, K. L., Nussbaum, A. M. & Wang, G. S. The pharmacologic and clinical effects of medical cannabis. Pharmacotherapy 33, 195–209, doi:10.1002/phar.1187 (2013).

18. Saito, V. M., Rezende, R. M. & Teixeira, A. L. Cannabinoid modulation of neuroinflammatory disorders. Curr. Neuropharmacol. 10, 159–166, doi:10.2174/157015912800604515 (2012).

19. Zurier, R. B. & Burstein, S. H. Cannabinoids, inflammation, and fibrosis. FASEB J. 30, 3682–3689, doi:10.1096/fj.201600646R (2016).

20. Cascio, M. G., Gauson, L. A., Stevenson, L. A., Ross, R. A. & Pertwee, R. G. Evidence that the plant cannabinoid cannabigerol is a highly potent alpha2-adrenoceptor agonist and moderately potent 5HT1A receptor antagonist. Br. J. Pharmacol. 159, 129–141, doi:10.1111/j.1476-5381.2009.00515.x (2010).

21. Silva, R. O. et al. Phytol, a diterpene alcohol, inhibits the inflammatory response by reducing cytokine production and oxidative stress. Fundam. Clin. Pharmacol. 28, 455–464, doi:10.1111/fcp.12049 (2014).

22. Baram, L. et al. The heterogeneity and complexity of Cannabis extracts as antitumor agents. Oncotarget 10, 4091–4106, doi:10.18632/oncotarget.26983 (2019).

23. Sledzinski, P., Zeyland, J., Slomski, R. & Nowak, A. The current state and future perspectives of cannabinoids in cancer biology. Cancer Med 7, 765–775, doi:10.1002/cam4.1312 (2018).

24. Ben-Shabat, S. et al. An entourage effect: inactive endogenous fatty acid glycerol esters enhance 2-arachidonoyl-glycerol cannabinoid activity. Eur. J. Pharmacol. 353, 23–31, doi:10.1016/s0014-2999(98)00392-6 (1998).

25. Morash, M. G. et al. Identification of minimum essential therapeutic mixtures from cannabis plant extracts by screening in cell and animal models of Parkinson’s disease. Front. Pharmacol. 13, 907579, doi:10.3389/fphar.2022.907579 (2022).

26. Andre, R. et al. The Entourage Effect in Cannabis Medicinal Products: A Comprehensive Review. Pharmaceuticals (Basel) 17, doi:10.3390/ph17111543 (2024).

27. Coles, M., Steiner-Lim, G. Z. & Karl, T. Therapeutic properties of multi-cannabinoid treatment strategies for Alzheimer’s disease. Front. Neurosci. 16, 962922, doi:10.3389/fnins.2022.962922 (2022).

28. Goldman, A. et al. Characterization of squamous esophageal cells resistant to bile acids at acidic pH: implication for Barrett’s esophagus pathogenesis. American journal of physiology. Gastrointestinal and liver physiology 300, G292–302, (2011). PMID: 21127259

29. Kimchi, E. T. et al. Progression of Barrett’s metaplasia to adenocarcinoma is associated with the suppression of the transcriptional programs of epidermal differentiation. Cancer Res. 65, 3146–3154, doi:10.1158/0008-5472.CAN-04-2490 (2005).

30. Stoner, G. D. et al. Establishment and characterization of SV40 T-antigen immortalized human esophageal epithelial cells. Cancer Res. 51, 365–371 (1991). PMID: 1703038

31. Nehra, D., Howell, P., Williams, C. P., Pye, J. K. & Beynon, J. Toxic bile acids in gastro-oesophageal reflux disease: influence of gastric acidity. Gut 44, 598–602, doi:10.1136/gut.44.5.598 (1999).

32. Payne, C. M., Bernstein, C., Dvorak, K. & Bernstein, H. Hydrophobic bile acids, genomic instability, Darwinian selection, and colon carcinogenesis. Clin Exp Gastroenterol 1, 19–47, doi:10.2147/ceg.s4343 (2008).

33. Goldman, A. et al. Protective effects of glycoursodeoxycholic acid in Barrett’s esophagus cells. Dis Esophagus 23, 83–93, doi:10.1111/j.1442-2050.2009.00993.x (2010).

34. Morales, P. & Reggio, P. H. An Update on Non-CB1, Non-CB2 Cannabinoid Related G-Protein-Coupled Receptors. Cannabis Cannabinoid Res 2, 265–273, doi:10.1089/can.2017.0036 (2017).

35. Zhao, P. & Abood, M. E. GPR55 and GPR35 and their relationship to cannabinoid and lysophospholipid receptors. Life Sci. 92, 453–457, doi:10.1016/j.lfs.2012.06.039 (2013).

36. Yin, H. et al. Lipid G protein-coupled receptor ligand identification using beta-arrestin PathHunter assay. J Biol Chem 284, 12328–12338, doi:10.1074/jbc.M806516200 (2009).

37. Lorenzen, E. & Sakmar, T. P. Receptor Structures for a Caldron of Cannabinoids. Cell 176, 409–411, doi:10.1016/j.cell.2019.01.012 (2019).

38. Tao, F., Zhang, Y. & Zhang, Z. The Role of Herbal Bioactive Components in Mitochondria Function and Cancer Therapy. Evid. Based Complement. Alternat. Med. 2019, 3868354, doi:10.1155/2019/3868354 (2019).

39. Bridgeman, M. B. & Abazia, D. T. Medicinal Cannabis: History, Pharmacology, And Implications for the Acute Care Setting. P T 42, 180–188 (2017). PMID: 28250701

40. Gertsch, J. Analytical and Pharmacological Challenges in Cannabis Research. Planta Med. 84, 213, doi:10.1055/s-0044-101051 (2018).

41. Eggertson, L. The challenges of medical cannabis research. Can. Nurse 113, 26–28 (2017). PMID: 29236415

42. Rodrigues, C. M., Fan, G., Wong, P. Y., Kren, B. T. & Steer, C. J. Ursodeoxycholic acid may inhibit deoxycholic acid-induced apoptosis by modulating mitochondrial transmembrane potential and reactive oxygen species production. Mol. Med. 4, 165–178 (1998). PMID: 9562975

43. Hebert-Chatelain, E. et al. A cannabinoid link between mitochondria and memory. Nature 539, 555–559, doi:10.1038/nature20127 (2016).

44. Bang, M. H. et al. Phytol, SSADH inhibitory diterpenoid of Lactuca sativa. Arch. Pharm. Res. 25, 643–646 (2002). DOI: 10.1007/BF02976937

