## Supplementary figures and images for "Optimal Cannabinoid-Terpene Combination Ratios Suppress Mutagenicity of Gastric Reflux in Normal and Metaplastic Esophageal Cells"

### Suppl. Fig 1

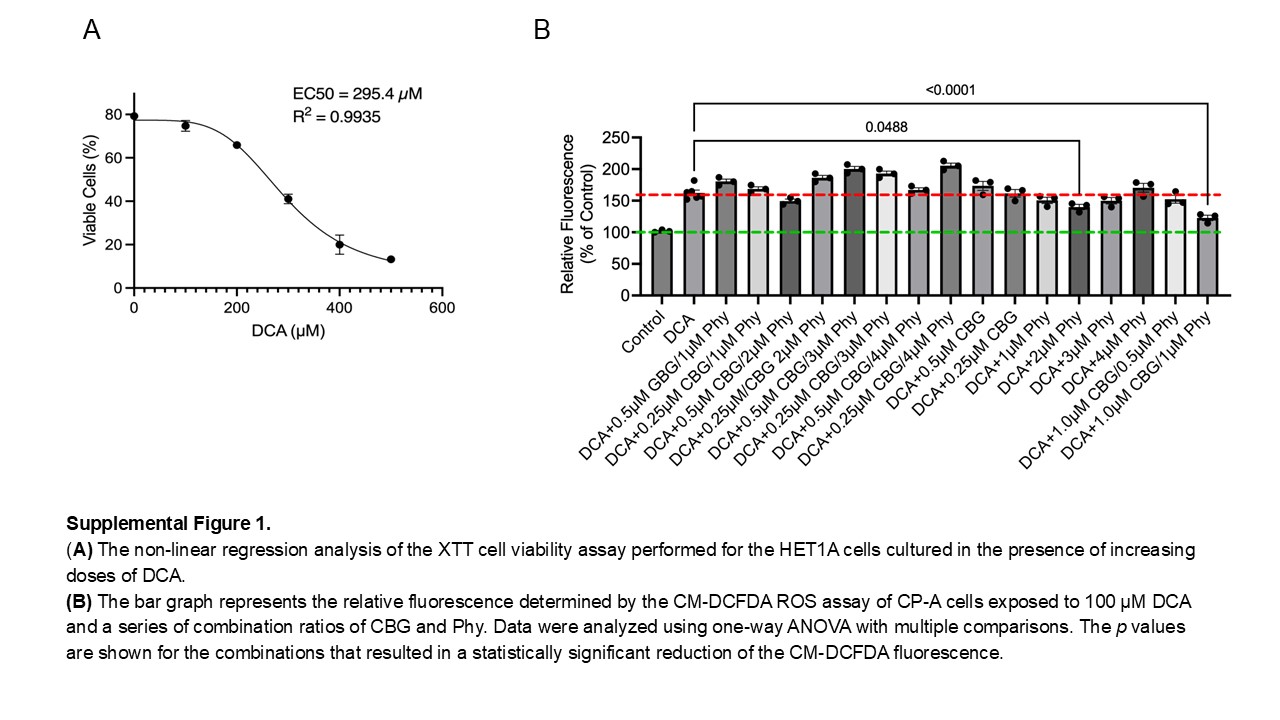

### Suppl. Fig 2

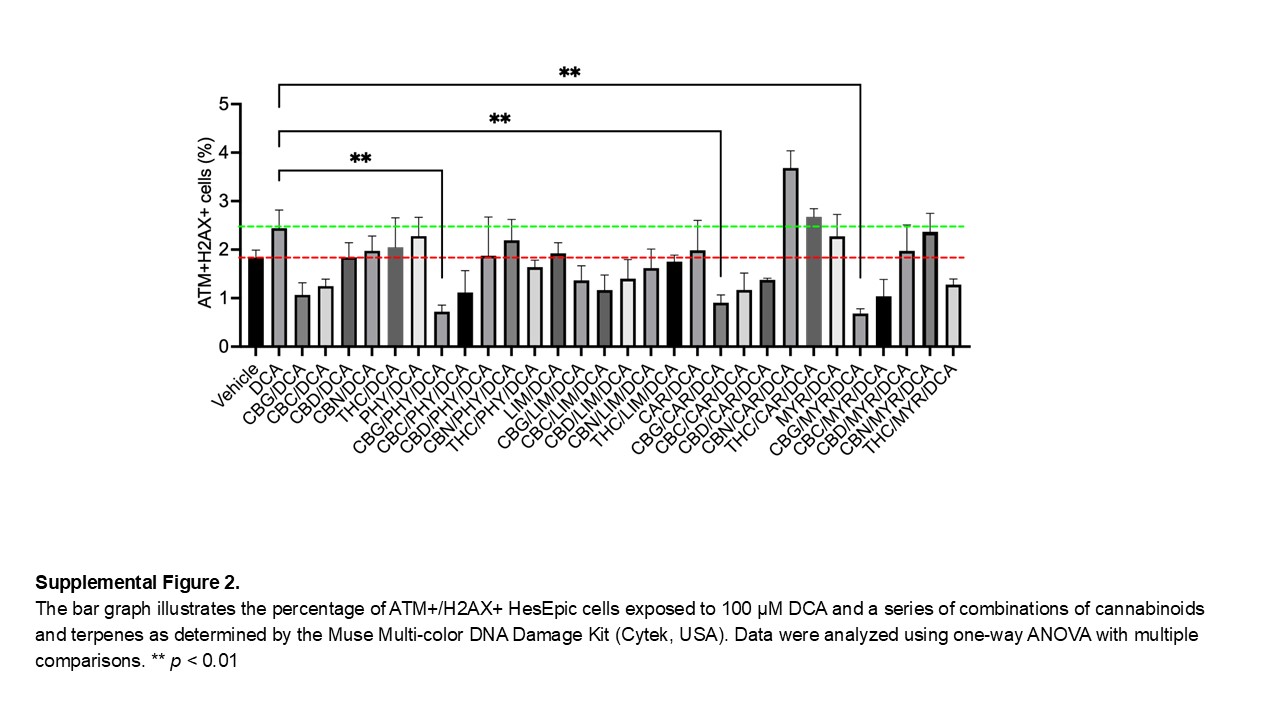
